# Developing and validation of a questionnaire for assessment of individual’s perceived risk of four major non-communicable diseases in Myanmar

**DOI:** 10.1101/2020.05.26.116202

**Authors:** Kyaw Swa Mya, Ko Ko Zaw, Khay Mar Mya

## Abstract

The adopting healthy life styles are greatly influenced by individual’s perceived risk of developing non-communicable diseases (NCDs). This study aimed to develop and validate a questionnaire that can assess the individual’s perceived risk of developing four major NCDs. Exploratory sequential mixed methods design was used. Qualitative part developed the question items pool by conducting two expert panels while quantitative part validated the questionnaire using both exploratory (EFA) and confirmatory factor analysis (CFA). Separate samples were used for EFA (n=150) and CFA (n=210). The participants were aged between 25-60 years of both sexes with no known history of NCDs. Face to face interview was conducted. Parallel analysis was done to decide the number of factors to be extracted. EFA was done using maximum likelihood method with Promax rotation to extract the underlying factors of perceived risk while CFA was done to assess the goodness of fit of proposed EFA Model using model fit indices. Based on literature search, 86-item questionnaire was firstly generated. During two expert panels, some overlapped items and items that did not represent the specific construct were removed. Experts made sure the content validity of developed 51-item questionnaire which was used to collect data from 360 participants. EFA revealed the five factors model with 22 high loading items which extracted 54% of total variance. CFA proved that hypothesized five factors model of 21-item questionnaire (one item was removed due to low loading) was satisfied with adequate psychometric properties and model fit indices (RMSEA=0.056, CFI=0.921, TLI=0.908, SRMR=0.063 & χ^2^/df=1.66). Developed 21-item questionnaire was shown to be valid and reliable to assess the perceived risk of developing NCDs among Myanmar population. Further research should be conducted to assess on the utility of the questionnaire in mismatch between risk perception and current risk and individualized counseling for behaviour change communication.

## Introduction

The World Health Organization (WHO) estimates that non-communicable diseases (NCDs) was responsible for 41 million people death in each year and accounts 71% of all deaths globally [1]. NCDs become the major public health problems for developing countries such as Myanmar and it has been recognized as a major challenge to achieve sustainable development goals. The WHO also states that 8.5 millions of lives are estimated to be lost due to NCDs in the South East Asia Region [2]. Myanmar, as one of 23 high burden countries with respect to NCDs [3] encountered a significant burden of NCDs and high potential to increase in exposure to risk factors associated with key NCDs in future [4]. The WHO 2^nd^ Global Status report on NCDs estimated that more than 50% of total deaths in Myanmar are accounted to NCDs and the probability of dying from one of the 4 major NCDs (Cardiovascular diseases (CVDs), Diabetes, Cancer and Chronic respiratory diseases) was about 24% in people aged 30 to 70 years. The report highlights a growing concern of several risk factors for NCDs including hypertension and overweight/obesity. It also pointed out that Myanmar had an increase in premature death from causes attributable to NCDs over the period between 1990 and 2010 [5]. According to the STEP survey in Myanmar (2014), among the people aged 40-64 years, 12% had already had one kind of CVDs or a high level (i.e., ≥30%) of 10-year CVDs risk. Nine out of 10 respondents had at least one NCDs risk factor and one out of five respondents had 3-5 risk factors in combination. These findings supported the national concern of growing several risk factors for NCDs in Myanmar [6].

NCDs are diseases that related to individual’s behavior. Most of the risk factors were modifiable. The potential to take necessary preventive measures for a particular disease was greatly influenced by individual’s perceived risk of developing these diseases [7]. Hence individual’s perception of developing disease or the belief of getting an adverse event among them is more important to adopt healthy lifestyle regardless of actual risk of developing these diseases [8]. For those at high risk, an accurate understanding of risk can realize patients to identify and adopt relevant lifestyle changes and follow the required preventive interventions that can lead to a better health-related quality of life [9–11]. For those at low or average risk, accurate risk perception can reduce anxiety and do not need to follow unnecessary sophisticated intervention [12].

Hence exploring the individual’s perception on developing NCDs using a standardized and validated tool specific to country context is necessary to combat the current increasing trend of NCDs risk factors in Myanmar. The best way to measure individual’s perception on developing NCDs is using a questionnaire since it is widely used by many researchers due to easy, cheap and its significant role in epidemiological data collection. Moreover, it is widely used as a research tool in public health surveys, especially in studies with large sample size, enabling statistical analysis that will give more power compared to other methods [13]. However, there are no standardized and validated questionnaires that can measure the perceived risk of developing NCDs in Myanmar. Hence, this study aims to develop and validate a questionnaire that can assess the individual’s perceived risk of developing four major NCDs within country specific socio-economic context.

## Materials and methods

The study used exploratory sequential mixed methods design and was conducted by 6 phases – 3 phases in qualitative approach and 3 phases in quantitative approach.

### Qualitative approach (Questionnaire development)

#### Phase 1. Conceptualization of constructs

This phase was started with reviewing the literature concerning with perceived risk based on health behavioral models. The study used the constructs of Health Belief Model (HBM) [14] – perceived susceptibility, perceived severity, perceived benefits, perceived barriers, perceived self-efficacy and perceived cues to action (behavioural change intention) to develop the questionnaire items pool for assessing the perceived risk of major NCDs since it was simple, effective and most popular behavior model.

#### Phase 2. Item generation (draft questions) and modification of questions by expert panel (Delphi method) to obtain satisfactory content validity

Initial item pool was generated from previous national and international researches, articles, conference papers by searching online, libraries and other available sources. Then two expert panels were conducted using Delphi method to ensure content validity of selected item pool. The experts were invited to take part voluntarily in this research project. The experts checked the items whether these items should be included or not in item pool and they also added item that should be included in item pool. Four point Likert scale was used for each selected item in the questionnaire to collect data from respondents.

#### Phase 3. Pilot testing to modify questionnaires to obtain satisfactory face validity

To ensure its comprehensibility and readability, pretest was done among 15 people aged 25 to 60 years of both sexes who had no known NCDs and were able to read and understand Burmese (Myanmar language) very well. Moreover, participants had been asked to respond to questions about clarity, content, appropriateness, and format of questionnaire items. Before final data collection, the questions were edited according to the participants’ suggestions to make sure translational validity (face validity).

### Quantitative approach (Questionnaire Validation)

#### Phase 4. Data collection and item purification by exploratory factor analysis (Questionnaire development)

A total of 360 participants, aged 25 to 60 years of both sexes without known NCDs were selected by consecutive sampling from outpatient department of Yangon General Hospital, North Okkalapa General Hospital and other teaching hospitals from Yangon region from September to December, 2019. The participants were patient’s attendants, workers and office staff who met with inclusion criteria. After getting informed consent, data was collected using pretested questionnaires at room or place where privacy was ensured. After data collection, the study provided health education pamphlets (Standardized Health Messages book, MOHS, Page no. 139, 140, 141) regarding to NCDs prevention to all participants [15]. These participants were randomly split into two groups – 150 participants for exploratory factor analysis (EFA) and 210 participants for confirmatory factor analysis (CFA).

The major objective of this phase was to reduce the number of items and evaluate the robustness of the intended items. Univariate analysis was done among 150 participants to assess item facility and item discrimination to ensure the selected items were appropriate for EFA. Items with reverse scoring were recoded to get conceptual direction of the construct. Whether items were answered in the same direction was examined using the facility index—approached extreme scores or had a low SD. To assess whether the sample size was adequate and the collected items were appropriate for EFA, Kaiser-Meyer-Olkin (KMO) measure of sampling adequacy and a Bartlett’s test of sphericity (Observed correlation matrix was not identical matrix) were assessed [16].

To determine the factorial structure of the questionnaire and which items together constituted a particular construct, an EFA—a widely used technique in exploring theoretical construct was used [17]. EFA was done using maximum likelihood factor extraction method with Promax rotation. Parallel analysis – the method compares the Eigenvalue obtained from the data matrix to the eigenvalues generated from a Monte-Carlo simulated matrix created from random data of the same size was done to determine the optimum number of factors to be extracted [18]. Parallel analysis scree plot was used to visualize the number of factors needed to be extracted. A number of iterations of EFA were carried out to constitute core items in each factor. Items were assessed in discriminating between participants’ responses to the questionnaire’s constructs i.e. latent factors (perceived susceptibility, perceived severity, perceived benefits, perceived barriers, perceived self-efficacy and perceived behavioural change intention). Discrimination of items was measured by inter-factors correlation and if the study found some factors were strongly correlated i.e. >0.7, EFA was rerun only with items of these factors to identify the item that correlated with both factors. The item correlated with both factors was removed to increase discriminant validity. The items were extracted based on not only factor loading but also interpretability of the factors. Items with low factor loading <0.40 and cross-loading with the difference below 0.2 were removed at each step of iteration [19].

#### Phase 5. Reliability of the questionnaire

Internal consistency (reliability) of each latent factor was tested for developed questionnaires by Cronbach’s alpha (α) coefficient and α ≥0.70 indicates good reliability [20]. The items that affect the reliability of latent factors were removed to get reliable factor. EFA was rerun excluding every item deleted for reliability reasons to get final EFA model.

#### Phase 6. Confirmatory factor analysis to check constructs validity (Questionnaire development)

To statistically confirm the EFA proposed perceived risk constructs on developing the NCDs, CFA was done using the remaining 210 participants’ data by assessing the convergent and discriminant validity of the constructs [21] and model fit measures using Structural Equation Modeling (SEM) technique. Model fit statistics such as root mean square error of approximation (RMSEA <0.08), comparative fit index (CFI >0.9), Tucker Lewis index (TLI>0.9), relative chi-square (χ^2^ /df <3), coefficient of determination (CD>0.95) and standardized root mean square residual (SRMR ≤0.08) were used to assess the model fitness [22]. Construct reliability was also assessed and reliability ≥0.7 indicates good reliability. All analyses were done using STATA (Version 15.1).

## Results

### Development of question items pool (Qualitative Phase)

Figure 1 shows the sequential development of final 21-item questionnaire from 86-item questions pool (Both questionnaire development and validation phase). Thorough literature review was done on PubMed, Google scholar to identify the items that can assess perceptions of non-communicable diseases. A total of 86-item questionnaire (S1 Table) was developed based on validated questionnaires that found in literature [13,23] and formulation of new items based on behavioral theory and Myanmar culture. These questionnaires were adapted appropriately to address Myanmar’s culture and tradition since the original items were not intended for assessing perceived risk on developing major NCDs among Myanmar population. Moreover, some items were formulated not only to accordance with the constructs of HBM but also to reflect country’s traditional beliefs and customs. Among 86-item questionnaire, 25 items were related to perceived susceptibility of disease, 9 items for perceived severity, 9 items for perceived benefit, 12 items for perceived barrier, 11 items for self-efficacy and 20 items for behavioral change intention.

**Figure 1.**
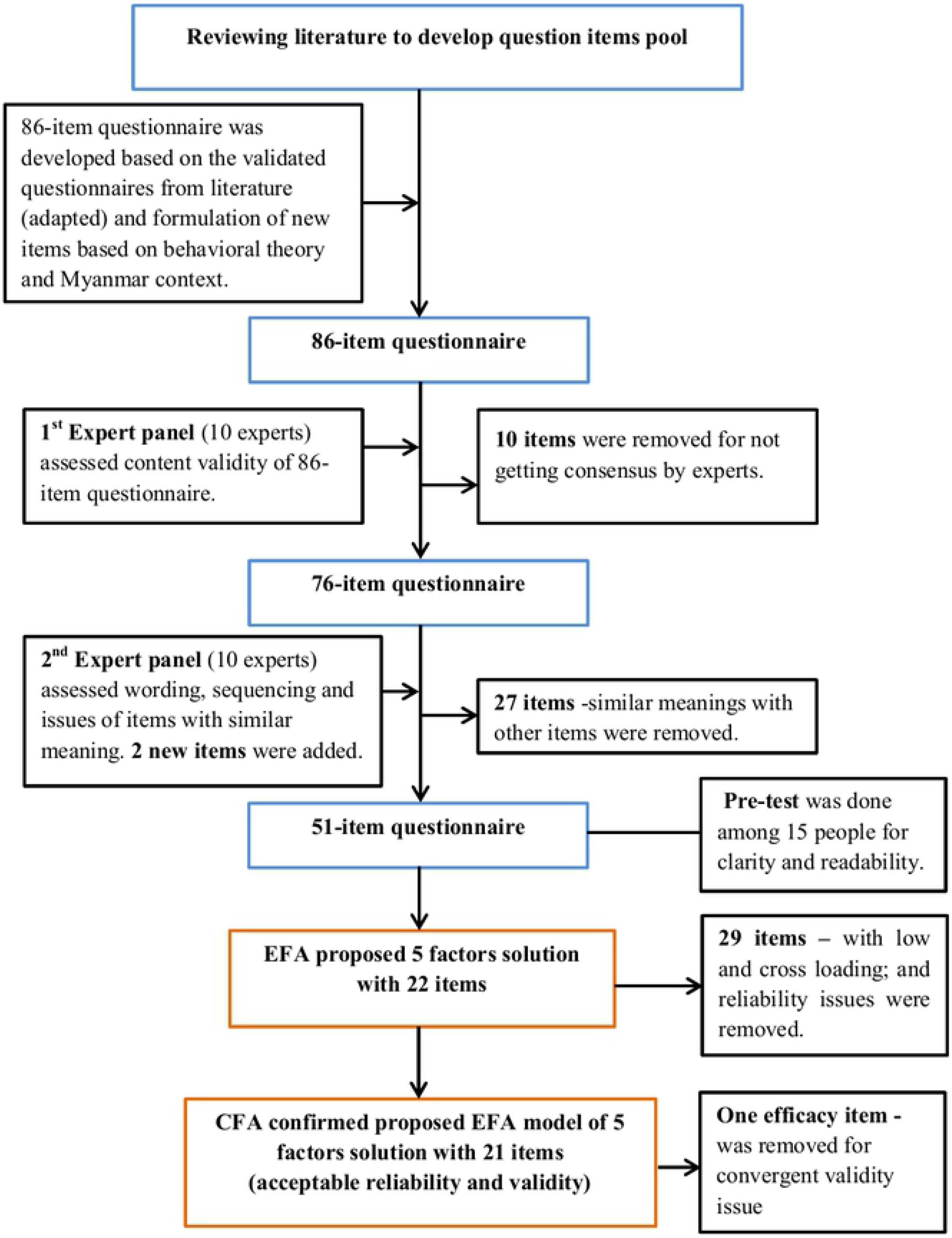
Questionnaire development and validation process.

To check the content validity, two expert panels were conducted with 10 experts including Clinician, Public Health specialists, Epidemiologists, Health policy expert, Social scientist, Demographer, Public health administrator, experienced researcher on NCDs. During 1^st^ expert panel, experts assessed the 86-item questionnaire using predefined 3 responses i.e. not at all representative, somewhat representative, or clearly representative. Among 86 items, 10 items (Item number 8, 9, 11, 13, 16, 20, 29, 32, 48 and 84) were removed for not getting consensus as clearly representative by more than 60% of experts. (S1 Table)

During 2^nd^ Delphi round, the experts noticed that some items group had similar meaning; therefore, these issues were resolved by taking consensus among experts to choose items that definitely reflect the perceived risk on developing major NCDs for specific construct. Among items of perceived susceptibility construct, five item groups (1, 4, 6, 7), (2, 21, 22, 23, 24, 25), (3, 17), (10, 14, 15, 16) and (18, 19) had similar meanings. Hence, experts agreed to select items (4, 6) from 1^st^ group, (22, 23, 24) from 2^nd^ group, (3) from 3^rd^ group, (10, 15) from 4^th^ group and (18) from last group. An expert also pointed out to remove item-15 since it was not concerned with susceptibility. Regarding to perceived severity items, two items (26, 33) had similar meaning so experts selected only item 26 to include in the final questionnaire. In perceived benefit items, experts agreed to select items 35 and 39 from two item groups (35, 38) and (36, 39). No similar issue was found in perceived barrier domain. Among self-efficacy items, item numbers (56, 60, 61) were similar. After taking consensus among experts, they agreed to retain only two items 60 and 61. They removed item 58 since this item was intended only to assess general preventive measures of NCDs and many items that assess specific preventive measures were already included in questionnaire. Regarding to behavioural change intention items, six item groups (67, 73, 76, 80), (68, 70), (69, 71, 79), (74, 78), (72, 75, 77, 81), (82, 83) had similar meanings, hence, they selected only items 67, 68, 71, 74, 75 and 82 from these groups. The experts also removed item 85 due to item itself was not related to perceived risk of NCDs. The items which are specific and not similar with other items in respective constructs were remained in the questionnaire. After 2^nd^ expert panel, 51-item questionnaire – 10 items in perceived susceptibility, 6 items in perceived severity, 7 items in perceived benefit, 11 items in perceived barrier, 9 items in self-efficacy and 8 items in behavioural change intention constructs, was developed and assured content validity by experts. Experts also revised and edited the wording of the items and sequencing of items to make sure understandability. Pretest was also done among 15 people aged 25-60 years, of both sexes and middle to graduate education level to assess the clarity and readability of developed 51-item questionnaire (S2 Table). Further modifications of wording of the items were done according to pilot testing results and suggestions not only from the participants but also from the interviewers to assure face validity.

### Questionnaire validation (Quantitative part)

To validate the developed questionnaire, the collected sample of 360 participants was randomly split into two separate samples. A sample of 150 participants was used for exploratory factor analysis and another sample of 210 participants was used for confirmatory factor analysis. Univariate analysis of the items was done and it was found that all the items’ means were ranged from 1.9 to 3.5 and their standard deviations were ranged from 0.5 to 1. The study also assesses the normality measure of the items and the results proved that there was no problem with skewness but kurtosis was existed among the items (S3 Table).

Before doing EFA, the study tested the two important assumptions of EFA i.e. Sampling Adequacy (KMO) to assess the adequacy of sample size and Bartlett’s test of sphericity to make sure the items were adequately correlated each other for EFA. KMO measure indicated the sampling adequacy since KMO value was greater than 0.7 (KMO=0.783). Bartlett’s test of sphericity test rejected the null hypothesis that correlation matrix was identical (P<0.001). Hence all assumptions were met and EFA analysis was done.

The study was based on health belief model to assess the perceived risk on developing non-communicable diseases, hence; questionnaire items were developed based on 6 constructs (latent factors) – perceived susceptibility, perceived severity, perceived benefit, perceived barrier, perceived self-efficacy and behavioural change intention. To decide the number of factor to be extracted from the EFA sample, parallel analysis which compares the Eigenvalue generated from the data matrix to the eigenvalues generated from a Monte-Carlo simulated matrix created from random data of the same size was done and the results were presented in Scree Plot (Figure 2). Parallel analysis proved that 5 factors (constructs) solution was the best for the EFA analysis since the simulated results cross the actual results between factor 5 and 6.

**Figure 2.**
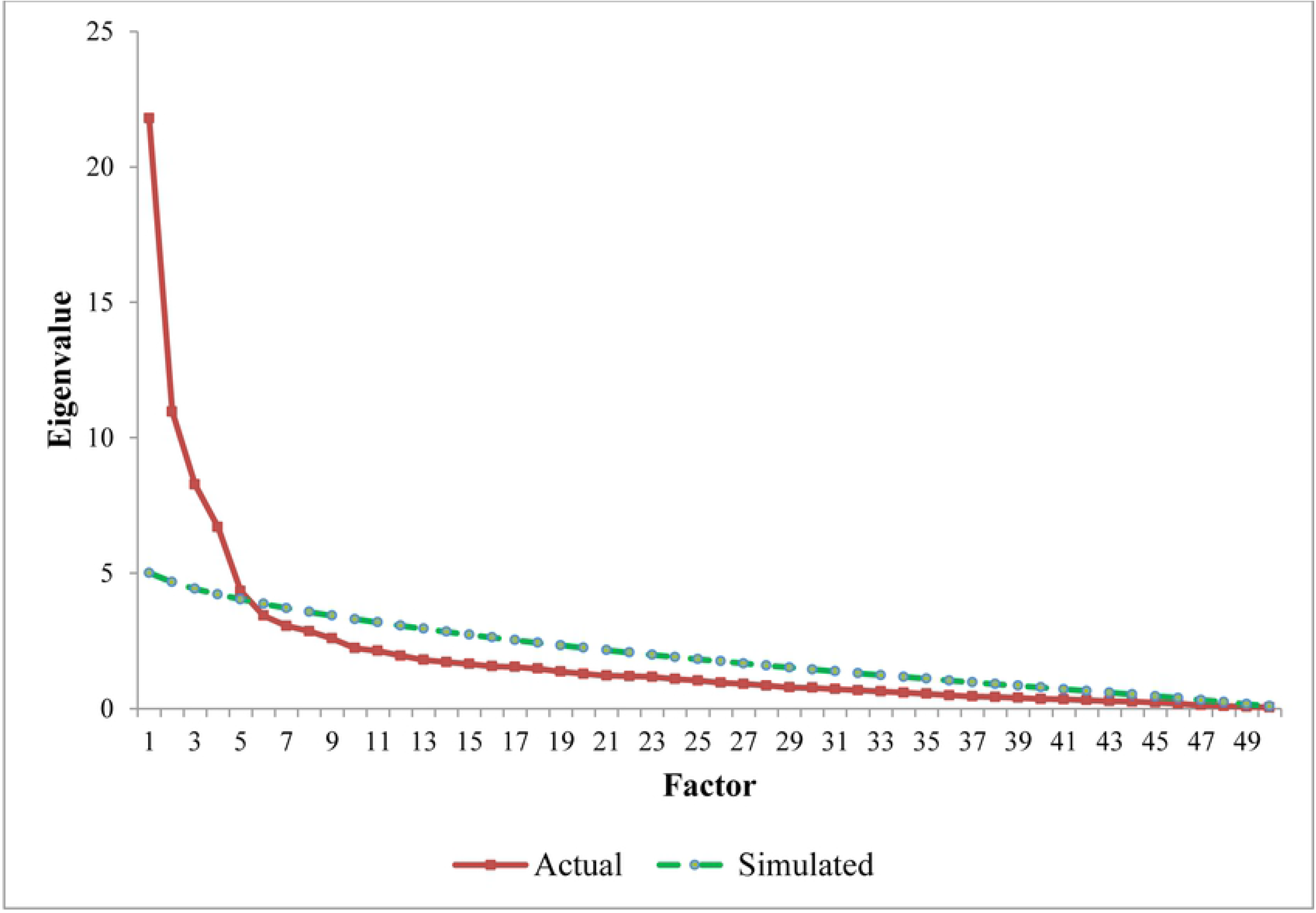
Parallel analysis Scree plot for decision on numbers of factors to be extracted.

### Exploratory factor analysis

EFA was done using maximum likelihood factor extraction method with Promax rotation with Kaiser Normalization. The first EFA output revealed that among 51 items, 10 items loaded in factor 1, nine items loaded in factor 2, nine items in factor 3, eight items in factor 4 and seven items in factor 5. Among these items, item sus_1 and sus_2 cross loaded in factor 3 and 5; and item intent_6 cross loaded in factor 1 and 4 (S4 Table – Pattern matrix 1). To get clean and theoretical meaningful results, the cross loading items were removed from EFA one after another and repeated EFA again. First, sus_2 was removed since the difference between two cross loadings was lowest and reanalyzed EFA again. Then sus_1 was removed and EFA was run again. During this analysis, previously unloaded item sus_4 was cross loaded between factor 2 and 5 with the difference loading 0.075 so this item was removed and reanalyzed again. After removal of intent_6, there was no more cross loading but two barrier items (bar_1, bar_2) were loaded together with behavioural change items and one behavioural change item (intent_8) was also loaded with barrier items. To get reasonable and theoretically interpretable constructs, these items were removed and rerun EFA again. After these item purification steps, eight self-efficacy items were loaded in factor 1, six benefit items and two severity items were loaded together in factor 2, eight barrier items loaded in factor 3, five behavioural change items loaded in factor 4 and; five susceptibility items loaded in factor 5. Factors were named according to loading items i.e. perceived self-efficacy (PerEffi) for factor 1, perceived benefit (PerBene) for factor 2, behavioural change intention (PerIntent) for factor 3, perceived susceptibility (PerSus) for factor 4 and perceived barrier (PerBar) for factor 5 (S4 Table – Pattern matrix 2).

To get reliable construct, reliability analysis was done for each factor using Cronbach’s alpha and some items that affected the reliability of factors were removed to increase the reliability. For PerSus factor, alpha value for five items was 0.774 and removing item sus_3 increased alpha to 0.792. For PerBene factor, alpha for eight items was only 0.670 and removal of item bene_1, seve_3 and seve_4 increased alpha to 0.831. Alpha value of PerBar was 0.757 using eight barrier items and that of PerEffi was 0.837 for eight efficacy items. PerIntent factor’s reliability was found to be 0.846 with five behavioural change intention items. Removing items to increase the reliability of factors affects the loading of factors so EFA was done again and again for removal of every item for reliability reason and the final EFA results were presented in Table 1. The main objective of this study was not only to develop the questionnaire but also to validate the questionnaire using confirmatory factor analysis (Structural Equation Modeling technique), hence, only the items with loading above >0.5 were included in the final model. The reliabilities of each latent factor were recalculated for the final model and the results were presented with factor loadings and percentage of variance extracted by each factor (Table 1).

**Table 1.** Factor loading results from exploratory factor analysis (22 items) and internal reliability of the factors.

| Item | Factor |  |  |  |  | Communalities |
| --- | --- | --- | --- | --- | --- | --- |
|  | PerEffi | PerBene | PerIntent | PerSus | PerBar |  |
| effi_7 | **.809** | -.124 | .055 | -.067 | .020 | .624 |
| effi_8 | **.742** | .170 | -.118 | -.105 | .039 | .578 |
| effi_9 | **.686** | -.069 | .141 | -.041 | -.050 | .535 |
| effi_2 | **.664** | -.072 | .055 | .074 | -.111 | .500 |
| effi_3 | **.558** | -.020 | .098 | .122 | -.022 | .371 |
| effi_6 | **.535** | .250 | -.137 | .069 | .054 | .363 |
| bene_3 | .024 | **.724** | .116 | -.051 | .140 | .669 |
| bene_2 | .051 | **.720** | .026 | -.081 | .208 | .617 |
| bene_7 | .107 | **.709** | -.177 | .210 | -.176 | .488 |
| bene_5 | -.088 | **.646** | .114 | -.014 | -.135 | .500 |
| bene_4 | -.053 | **.626** | .180 | -.069 | -.081 | .554 |
| intent_3 | .109 | -.085 | **.862** | .042 | .026 | .711 |
| intent_2 | .163 | .021 | **.750** | .009 | .063 | .664 |
| intent_4 | -.132 | .142 | **.689** | .079 | -.143 | .589 |
| intent_1 | -.046 | .143 | **.658** | -.053 | .048 | .550 |
| sus_5 | .023 | -.053 | .018 | **.745** | .047 | .570 |
| sus_6 | -.096 | -.060 | .004 | **.736** | .056 | .569 |
| sus_8 | .105 | .038 | -.093 | **.691** | .036 | .523 |
| sus_10 | -.017 | .074 | .153 | **.640** | -.027 | .413 |
| bar_4 | .028 | -.070 | .001 | -.015 | **.768** | .577 |
| bar_7 | -.141 | .089 | .140 | .072 | **.619** | .458 |
| bar_5 | .024 | -.029 | -.166 | .081 | **.553** | .383 |
| **Cronbach's $\alpha$** | .834 | .831 | .854 | .792 | **.683** |  |
| **% of variance** | 23.405 | 11.004 | 9.994 | 5.258 | 4.005 |  |
| **Cumulative %** | 23.405 | 34.409 | 44.403 | 49.662 | 53.667 |  |

PerEffi= Perceive self-efficacy, PerBene=Perceived benefit, PerIntent= Perceived behavioural change intention, PerSus= Perceived susceptibility, PerBar= Perceived barrier, CR= Construct reliability

Table 1 describes the factor loading results of final exploratory factor analysis and it was shown that 22 items loaded strongly to 5 factors. Six efficacy items significantly loaded to PerEffi factor which accounted for 23.4% of total variance with reliability alpha 0.834. Five benefit items highly loaded to PerBene factor and accounted 11% of total variance with alpha 0.831. Four behavioural change intention items were loaded to PerIntent factor with reliability alpha 0.854 and accounted for 9.9% of total variance. Four susceptibility items loaded strongly to PerSus factor which accounted 5.3% of variance of data with reliability alpha 0.792. Only 3 barrier items loaded to Factor PerBar with 0.683 and this factor accounted 4% of total variance. More than 50% of total variance was extracted by 5 factor solution of EFA. Average factor loading of each factor was greater than or equal to 0.65 and this finding point out that convergent validity of each factor was satisfactory. Regarding to reliability, all the factors’ reliability exceed 0.7 except PerBar factor which has reliability nearly 0.7. These findings revealed that reliability of all factors were satisfied to conduct CFA for validation process of 22 items questionnaire developed by EFA.

Factor correlation matrix of final exploratory factor analysis was described in Table 2. It was found that both negative and positive correlation among 5 factors. The Highest negative correlation was found between PerEffi and PerBar (−0.249) and the lowest between PerSus and PerBene (−0.053). Highest positive correlation was found between PerBene and PerIntent (0.577) and lowest positive correlation between PerBene and PerBar (0.025). All these correlation coefficients were less than 0.7 which was the upper limit that determine the discriminant validity issue; hence, the factors derived from EFA revealed the adequate discriminant validity among the extracted factors.

**Table 2.** Factor correlation matrix of EFA.

| **Factor** | **PerEffi** | **PerBene** | **PerIntent** | **PerSus** | **PerBar** |
| --- | --- | --- | --- | --- | --- |
| PerEffi | 1.000 |  |  |  |  |
| PerBene | .280 | 1.000 |  |  |  |
| PerIntent | .290 | .577 | 1.000 |  |  |
| PerSus | .064 | -.053 | -.140 | 1.000 |  |
| PerBar | -.249 | .025 | -.159 | .136 | 1.000 |

Before CFA, Harman’s single factor test was done to assess the common method bias which is a systematic response bias and can occur when a single data collection method was used and that will either inflate or deflate response. This test uses the maximum likelihood method and forced to extract only one factor whether to assess a single factor contributes more than 50% of total variance. It was found that only 25.5% was accounted by one factor and this means that there was no problem with common method bias in this study (S5 Table).

### Confirmatory Factor Analysis

EFA provided 22 items questionnaire for underlying five factors that influenced the perceived risk on developing non-communicable diseases among adults. These 5 factors have sufficient convergent validity and discriminant validity to conduct the CFA. Moreover, all the factors satisfied with adequate reliability. Whether EFA proposed 5 factors with 22 items questionnaire model can be used to assess the perceived risk on developing non-communicable diseases at population level, CFA was done using different sample of 210 participants.

CFA initial model was run with 22 items that hypothesized by EFA and it was found that effi_6 item was loaded to PerEffi factor with low loading (0.35). Hence CFA final model was run with only 21 items (dropping effi_6 item). The results of CFA final model (NCD-PR5-21) were presented with SEM diagram in Figure 3. Now all the loadings were ranged from minimum 0.5 to maximum 0.8 and all correlation between the latent factors were less than 0.7.

**Figure 3.**
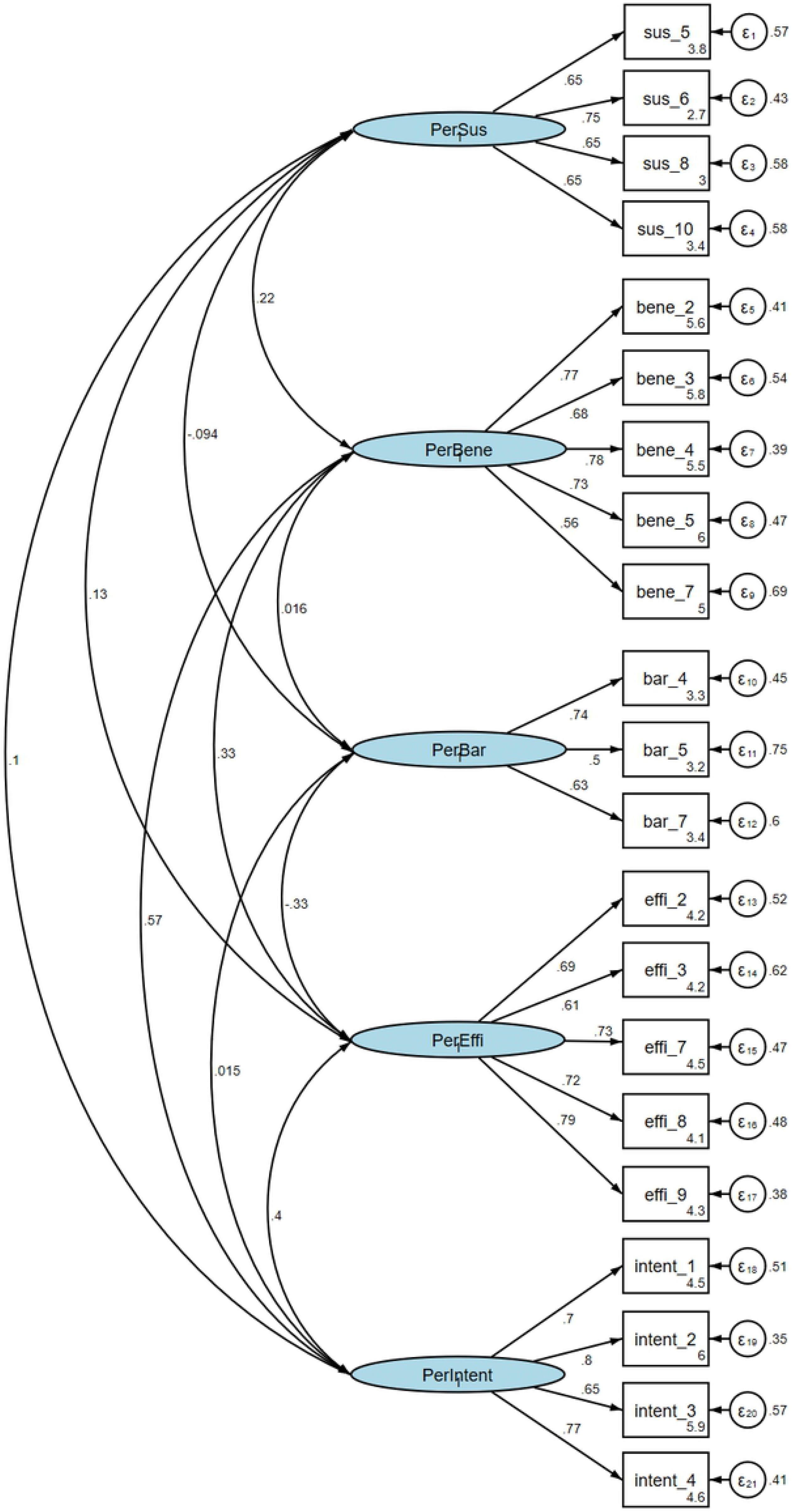
CFA final model (NCD-PR5-21) for perceived risk on developing non-communicable diseases.

Tables 3 describes the convergent validity, discriminant validity and construct reliability of CFA final model after removing low loading item i.e. effi_6. Regarding to construct reliability, all factors’ reliability were above the 0.7 except for PerBar factor whose reliability was 0.662. Average variance extracted (AVE) value of PerBene, PerEffi and PerIntent factors were above 0.5 while that of PerSus and PerBar were below 0.5. Although two factors i.e. PerSus and PerBar had AVE value less than 0.5, factors specific items loadings were acceptable for convergent validity since there was not items with loading below 0.5 (See in Figure 3). All square correlation values were lower than AVE values of their respective factors; hence, there was no issue for discriminant validity for CFA final model. Regarding to model fit indices of CFA final model, the findings illustrated good model-data-fit i.e. RMSEA <0.08, PCLOSE>0.05, relative chi-square <3, CFI and TLI >0.9, SRMR<0.08 and CD>0.95. Hence, CFA proved that perceived risk on developing NCDs had underlying 5 latent factors and can be assessed using 21-item questionnaire (NCD-PR5-21) at population level (See final questionnaire in S6 Table).

**Table 3.** Convergent validity, discriminant validity, reliability and model fit indices of CFA final model with 5 latent factors.

| Squared correlations (SC) among latent variables |  |  |  |  |  |  |  |  |
| --- | --- | --- | --- | --- | --- | --- | --- | --- |
|  | CR | AVE | PerSus | PerBene | PerBar | PerEffi | PerIntent |  |
| PerSus | 0.773 | 0.459 | (0.459) |  |  |  |  |  |
| PerBene | 0.829 | 0.502 | 0.048 | (0.502) |  |  |  |  |
| PerBar | 0.662 | 0.401 | 0.009 | 0.000 | (0.401) |  |  |  |
| PerEffi | 0.832 | 0.504 | 0.018 | 0.112 | 0.109 | (0.504) |  |  |
| PerIntent | 0.820 | 0.539 | 0.011 | 0.32 | 0.000 | 0.163 | (0.539) |  |
| Model Fit indices | | RMSEA | PCLOSE | $\chi^2/df$ | CFI | TLI | SRMR | CD |
|  |  | 0.056 | 0.0179 | 1.66 | 0.921 | 0.908 | 0.063 | 1.0 |

Note: when AVE values ≥ SC values there is no problem with discriminant validity, when AVE values ≥ 0.5 there is no problem with convergent validity, CR = Construct reliability ( ≥ 0.7 is good reliability)

## Discussion

This study started with qualitative approach to identify 86 items that reflect the underlying dimensions of popular health behavioural model – HBM regarding to the perception on NCDs. Then 51-item questionnaire was developed and satisfied content validity by conducting two rounds of expert panels. In quantitative approach, both EFA and CFA were done using the separate samples. Among predefined HBM constructs – perceived susceptibility, severity, benefit, barrier, efficacy, behavioural change intention, items of perceived severity factor were dropped during EFA and provided the hypothesis of the five factors solution with 22-item questionnaire. CFA also confirmed the hypothesis of five factors model of perceived risk on developing NCDs with 21 items (NCD-PR5-21) (one efficacy item was removed to increase convergent validity) with satisfactory level of psychometric properties i.e. acceptable convergent validity, discriminant validity and reliability; and model fit indices.

Factor analysis was a statistical method which can measure latent factors – measures that cannot be measure directly such as satisfaction, perception and burnout. Three main purposes of using it were: (1) to conceptualize the underlying components of a set of variables; (2) to develop psychometric tool based on theory; (3) to reduce the a set of variables to one or manageable numbers while minimizing the loss of information as in multicollinearity situation. The use of factor analysis in this study was to develop and validate the questionnaire assessing the perceived risk on developing NCDs. Factor analysis includes two different components – EFA and CFA. EFA was usually assumed as data driven approach since there was no a priori theory and it determines the underlying constructs based on correlation among the observed variables while CFA was theory based approach in which hypothesized model was tested with the collected data to prove that there was acceptable model fitness [24]. In our study, EFA first explore the relationship of the items to provide the hypothesized underlying latent constructs of perceived risk on NCDs using minimal number of items and then CFA confirmed the hypothesized model was good fit to assess the individual’s perceived risk using two different samples. Conducting both EFA and CFA in one sample was unacceptable since no new information could be obtained. Our study used standard psychometric tool development methodology conducting both EFA and CFA with different samples.

The widely used factor extraction techniques for EFA were Principal component analysis (PCA) and Principal axis factoring analysis (PAF), Maximum likelihood method, unweighted least squares and generalized least squares. All these methods except PCA extract factors based on communalities i.e. shared/common variance excluding unique variance while PCA used total variance without decomposing common or unique variance [17]. Moreover, PCA is mainly used for data reduction; in contrast, other methods intend to extract the underlying latent factors and parameter estimation. Hence, we used the maximum likelihood method instead of using PCA and PAF in order to maximize the likelihood of reproducing the population correlation matrix [25,26] since the study aimed to confirm EFA proposed model using CFA in which maximum likelihood method was used for parameter estimation. Decisions on how many factors should be extracted was also a major concern in factor analysis since both underfactoring and overfactoring cause problems i.e. difficult to interpret due to substantial error in underfactoring while developing unrealistic and complex theories in overfactoring. To avoid these problems, we used the parallel analysis method instead of using Kaiser Criterion and scree plot. It compares the eigenvalues produced be the real data with the eigenvalues estimated from Monte-Carlo simulated matrix created from random data [22,25,26] and the number of factors above the intersection point should be extracted. In our study, the simulated results cross the actual results between factor 5 and 6, hence, we extracted 5 factors from the data. Moreover, to prevent underfactoring or overfactoring, the study assessed not only 5 factors solution but also 4 and 6 factors solution. The findings of 4 and 6 factors solutions were not good enough for theoretical interpretation and some items were cross loaded with more than one factor (See detail in S7 Table). The study also used Promax rotation one of the oblique rotation method which allowed to correlate the extracted factors each other instead of using orthogonal rotation which does not allow to correlate among the factors since we belief the underlying factors of perceived risk were somewhat correlated each other and EFA results also described the significant correlation between some factors.

During EFA, the items that assessed the perceived severity of NCDs were dropped for many reasons. First, the parallel analysis provided the evidence to extract 5 factors solution; hence, we forced to 5 factors model and some severity items were loaded together with the benefit items and some items were low loading i.e. <0.4. Second, when we ran step by step EFA for item purification, these items were cross loading with other items, that’s why some of them were remove. Third, to get reliable construct, we assessed the internal consistency of constructs and some of these items needed to remove to increase the reliability of underlying constructs. These severity items were not able to strongly correlate with each other to form a factor like other items (S4 Table & S7 Table). This might be due to the fact that the participants were the one who had no known NCDs; hence they failed to perceive the severity of NCDs or the developed items had lack of intrinsic ability to capture their perception regarding to severity of diseases due to bias wording or ambiguous wording.

Regarding to model fit of CFA, there are many model fit indices in structural equation modeling methods i.e. absolute fit indices (χ^2^, relative χ^2^, GFI, SRMR), parsimonious fit indices (RMSEA, AIC, AGFI) and incremental fit indices (CFI, TLI) [27,28]. All these indices have some limitations depending on assumptions, sample size and model complexity. For example, chi-square value greatly influenced on sample size i.e. the theoretically plausible model can be rejected in case of sample size increases with constant number of degree of freedom. Hence some of the articles report relative chi-square which is calculated by dividing chi-square value by degree of freedom. The results of less than 3 indicate the good or acceptable model fit. Goodness of fit index (GFI) and adjusted goodness of fit index (AGFI) were used as model comparison but these indices depend on sample size and more likely to underestimate model fitness [29]. Many studies [27–29] recommended to use RMSEA, CFI and TLI since these indices are sensitive to model misspecification and less depend on sample size. In our study, we used relative chi-square, RMSEA, CFI, TFI, SRMR and CD. All these indices were at acceptable level of model fit and indicate the observed covariance matrix in CFA was identical with the proposed covariance matrix in EFA. Among five latent factors, only one factor has reliability less than minimum acceptable level but its reliability was nearly 0.7 i.e. 0.662 and it was acceptable. Moreover there were also convergent validity issues for PerSus and PerBar factors since their respective AVEs were below 0.5. However, all factors’ loading ranged from 0.5 to 0.8 and no item loadings were below 0.5 which indicates the convergent validities of these two factors were also acceptable [30].

### Strengths and limitation of the study

To the best of our knowledge, this study was the first study that developed and validated the questionnaire assessing the individual’s perceived risk on developing NCDs in Myanmar. The study used the standard psychometric tool development methodology i.e. designing, developing and testing measures of psychological constructs using mixed methods design and two separate samples for factor analysis. However, some items had low loading (<0.65) in CFA and perceived barrier factor had low reliability (0.662) pointed out that NCD-PR5-21 had limitation and should also be tested in other population with larger sample size to make sure the generalizability of the findings.

## Conclusion

Developed 21-item questionnaire (NCD-PR5-21) for 5 underlying factors – perceived susceptibility, benefit, barrier, efficacy, behavioural change intention was satisfied with acceptable level of validity and reliability, hence, it can be used to assess individual’s perceived risk on major NCDs among Myanmar population. Further research should be conducted not only to confirm reliability and validity but also to reproduce the factors determined the perceived risk on NCDs with an independent, larger sample to generalize the study’s findings. The questionnaire should also be tested on the utility of the questionnaire in mismatch between risk perception and current risk and individualized counseling for behaviour change communication.

## Acknowledgments

I want to show my gratitude to members of expert panel who gave great inputs in doing Delphi method in qualitative parts of this study. Without their academic inputs, the research could not be finished successfully. I would like to thank Chairman and members of Implementation Research (IR) grant of Ministry of Health and Sports, Myanmar for providing the funding support for this research. Last but not least, I would like to express my special thanks to all Master of Hospital Administration students (2019) and three Master of Public Health students (2019) – Dr. Tin Aung Cho, Dr. Zaw Phyo Aung and Dr. Paing Khant for their active participation in data collection and respondents of this study for their enthusiastic involvement in answering the survey questions.

## Supporting information

**S1 Table. Items pool for developing questionnaires for perceived risk on NCDs**

**S2 Table. Development of 51-item questionnaire**

**S3 Table. Univariate analysis of items that included in EFA analysis**

**S4 Table. Exploratory factor analysis – Initial step & Factor naming step**

**S5 Table. Harman’s single factor test to check for common method bias**

**S6 Table. Validated final 21-item questionnaire (NCD-PR5-21)**

**S7 Table. Exploratory factor analysis for six factor solution and four factor solution**

